# A Nonlinear Biomechanical Model for Prognostic Analysis of Clavicle Fractures

**DOI:** 10.64898/2026.04.06.716697

**Authors:** Yiyan Chen

## Abstract

Clavicle fractures often exhibit markedly different clinical outcomes: some patients recover acceptable function despite shortening or displacement, whereas others with apparently similar deformity develop persistent pain, functional loss, or poor healing. To explain this distinction, we propose a minimal nonlinear mechanical model for prognostic analysis of clavicle fractures. The model describes the interaction between fracture-related shortening and compensatory shoulder-girdle posture through a reduced equilibrium equation incorporating stiffness, geometric nonlinearity, and shortening-posture coupling.

Within this framework, we analyze equilibrium branches, local stability, and the emergence of critical thresholds. We show that post-fracture destabilization can be interpreted as a fold bifurcation, while more complex parameter dependence gives rise to cusp-type structures and multistability. These bifurcation mechanisms provide a mathematical explanation for sudden deterioration after injury or treatment, as well as for strong inter-individual variability.

We further introduce an optimization principle based on a utility functional to guide treatment planning. The analysis predicts that the optimal safe correction should lie strictly below the bifurcation threshold, thereby generating a natural safety margin. Although the model is simplified and has not yet been calibrated against patient data, it nevertheless provides a theoretical framework for understanding why fracture prognosis may deteriorate abruptly near critical mechanical conditions and offers a dynamical-systems interpretation of empirical treatment thresholds used in clinical practice.

## 1. Introduction

Clavicle fracture is one of the most common injuries of the shoulder girdle, with midshaft clavicle fractures being especially frequent. Yet clinical outcomes vary substantially. Under conservative treatment, some fractures heal uneventfully without functional sequelae, whereas other cases with apparently similar degrees of shortening or displacement lead to chronic pain, altered shoulder mechanics, restricted motion, or even nonunion. In practice, treatment decisions are often made empirically, based on whether shortening, displacement, comminution, or functional impairment appears severe enough to justify surgery. Although this threshold-like phenomenon is widely recognized, its underlying mechanism remains insufficiently understood.

This raises a natural question: why can some fractures remain mechanically tolerated, whereas others of comparable apparent severity lead to pronounced functional deterioration and poor healing? One possible explanation is that prognosis depends on fracture severity in a nonlinear rather than linear way. The shoulder girdle is not a simple rigid structure but a coupled mechanical system in which fracture-related shortening interacts with compensatory posture, joint loading, and soft-tissue response. From this perspective, a small additional amount of shortening may do more than increase risk gradually; it may push the system across a critical boundary beyond which the mechanically stable configuration is lost.

This viewpoint is consistent with a broad class of nonlinear systems in which equilibria branch and change stability. Such abrupt transitions are classically described by bifurcation theory. Motivated by this perspective, the aim of the present paper is to propose a minimal nonlinear mechanical model for clavicle-fracture prognosis. Rather than constructing a high-fidelity anatomical simulation, we introduce a reduced model intended to capture the essential mechanism behind threshold-like deterioration, inter-individual variability, and the existence of mechanically meaningful safety margins.

In the model, a geometric control parameter represents fracture-related shortening, an effective state variable represents compensatory shoulder-girdle posture, and nonlinear coupling terms encode stiffness, geometry, and baseline posture. We first analyze the equilibrium structure of the system and derive a criterion for local stability. We then show that loss of stability can be interpreted as a fold bifurcation of equilibrium branches. Clinically, this means that once shortening exceeds a threshold, mechanically stable compensation may disappear, leading to pain or restricted motion. We further show that when an additional baseline parameter varies, the fold set develops into a cusp-type geometry, giving rise to multistability and strong dependence on patient-specific conditions.

The second objective of this work is to connect the bifurcation structure with treatment planning. Clinical decision-making is almost never based on stability alone; in practice, it balances geometric correction against joint loading, muscular compensation, and proximity to instability thresholds. To formalize this idea, we introduce a utility-based optimization principle on the stable equilibrium branch. The resulting analysis predicts that the optimal safe correction should lie strictly below the bifurcation threshold and that a nonzero safety margin emerges naturally from the competition between benefit and fragility. In this sense, empirical treatment thresholds need not be interpreted merely as heuristic cutoffs; they may also reflect an underlying boundary of mechanical instability.

## 2. Model Formulation

### 2.1 State variable and control parameter

We assume that the degree of shoulder-girdle compensation after clavicle fracture can be represented by a scalar variable *ϕ*, with larger values corresponding to stronger compensatory adjustment. This variable is intended to summarize the combined effects of acromioclavicular motion, sternoclavicular motion, scapular adaptation, and soft-tissue response. In reality, such compensation is high-dimensional, but for the present preliminary theoretical study we reduce it to a single effective variable.

We denote by δ ≥ 0 a geometric control parameter representing fracture-related clavicular shortening. Here*δ* is not identified with any single imaging measurement. Rather, it is an abstract structural measure of the mechanical alteration induced by shortening, including both physical shortening and functionally equivalent geometric changes. This distinction is important because clinical measurements of “shortening” depend on imaging modality and measurement protocol.

A third parameter, *ϕ*_0_, represents the pre-fracture baseline state of the shoulder girdle and encodes patient-specific differences such as resting posture, ligamentous structure, joint configuration, muscular balance, and habitual movement patterns. Mathematically, *ϕ*_0_ introduces asymmetry and helps organize multistable behavior.

### 2.2 Reduced equilibrium equation

We describe mechanical equilibrium by the scalar equation

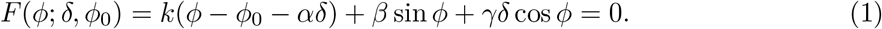

The first term, *k*(*ϕ*− *ϕ*_0_− *αδ*), represents a linear restorative tendency. Here *k >* 0 is an effective stiffness parameter, and *αδ* describes the shift in preferred configuration induced by fracture shortening. The parameter *α* measures how strongly shortening changes the mechanically preferred state.

The second term, *β* sin *ϕ*, represents intrinsic geometric nonlinearity associated with ligaments, joint capsule effects, and other restoring mechanisms that cannot be captured adequately by a purely quadratic energy. For small *ϕ*, it behaves approximately linearly, but for larger displacements it generates curved branches and bifurcation phenomena.

The third term, *γδ* cos *ϕ*, describes a shortening-dependent coupling between geometry and posture. Its role is to express the idea that shortening does not merely translate the preferred state linearly; it also changes how posture feeds back into equilibrium. This term is one of the principal sources of the model’s nontrivial bifurcation structure.

At this stage, these parameters are structural rather than directly calibrated clinical quantities. The purpose of the model is not yet quantitative prediction for individual patients, but rather identification of the competing mechanical tendencies that can generate threshold-like deterioration.

### 2.3 Energy formulation

Equation (1) can be interpreted as the stationarity condition of a reduced scalar energy:

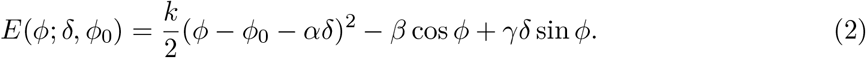

Indeed,

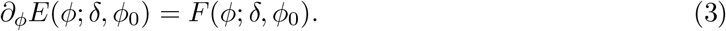

Thus, equilibria are precisely the critical points of the energy. In this representation, local minima correspond to locally stable equilibria, whereas local maxima correspond to unstable equilibria. The energy viewpoint is especially useful because it makes multistability intuitive: qualitatively different prognostic states correspond to distinct local minima of the same reduced energy landscape.

**Figure 1:**
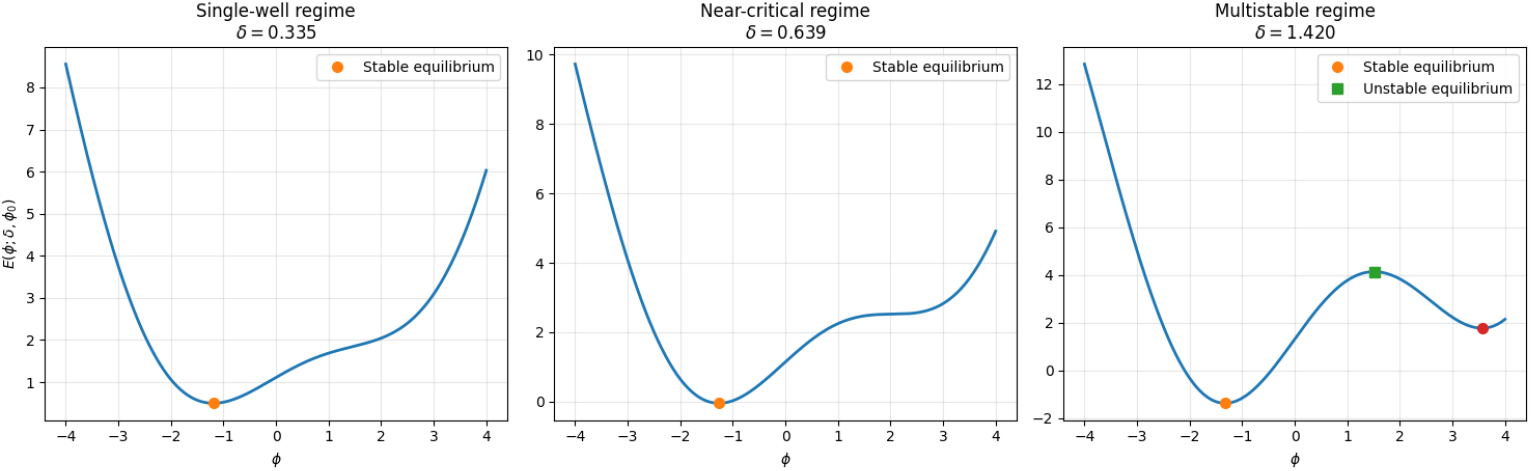
Representative energy landscapes of the reduced model, illustrating a monostable regime, a near-critical regime, and a multistable regime.

### 2.4 Stability criterion

Differentiating equation (1) with respect to *ϕ*, we obtain

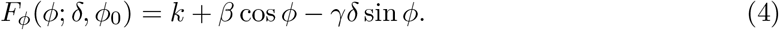

A given equilibrium *ϕ*^*^ is locally stable if

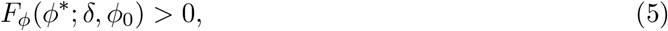

and unstable if

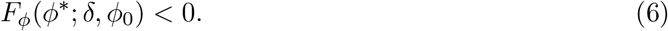

The critical case *F*_*ϕ*_ = 0 marks the loss of local invertibility of the equilibrium branch and will later be identified with the fold boundary. Mechanically, *F*_*ϕ*_ may be interpreted as an effective local stiffness margin: when it is comfortably positive, the equilibrium is robust; when it approaches zero, the system becomes fragile and highly sensitive to small perturbations.

## 3. Equilibrium Branches and Stability

The purpose of this section is threefold. First, we show that away from degenerate points the equilibrium branch can locally be expressed as a smooth function of the shortening parameter *δ*. Second, we quantify the sensitivity of this branch to shortening. Third, we show that the same quantity controlling branch sensitivity also controls local stability, thereby revealing a mechanism of pre-critical fragility.

### 3.1 Reference equilibrium at zero shortening

We begin with the reference problem*δ* = 0, for which the equilibrium equation reduces to

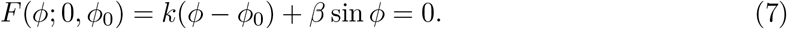

Let 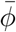 be a solution, so that

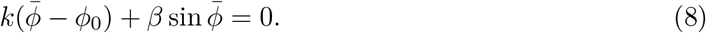

This gives the baseline equilibrium configuration in the absence of fracture-related shortening.

In general, 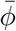 need not coincide exactly with *ϕ*_0_, since the nonlinear geometric term may shift the true equilibrium away from the nominal baseline parameter.

Now compute

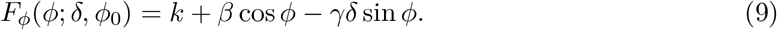

At the reference point,

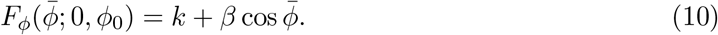

If

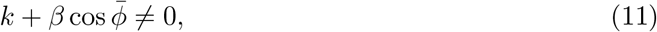

then the implicit function theorem implies the existence of a unique smooth local equilibrium branch *ϕ* = *ϕ*(*δ*) with 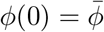.

#### Proposition 3.1

(Local equilibrium branch). *Suppose* 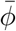 *satisfies (8)*. 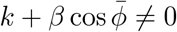, *then in some neighborhood ofδ* = 0, *the equilibrium equation F* (*ϕ*;δ, *ϕ*_0_) = 0 *defines a unique smooth branch*

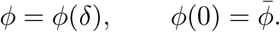

### 3.2 First-order sensitivity with respect to shortening

Differentiating the equilibrium identity

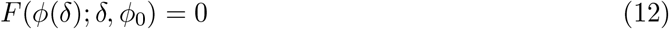

with respect to *δ*, we obtain

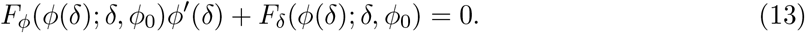

Since

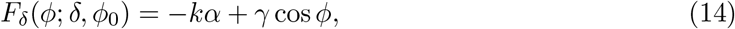

it follows that

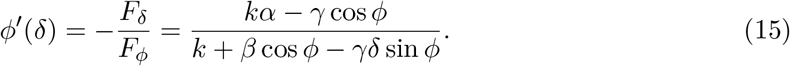

In particular,

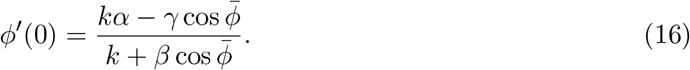

This formula has a direct mechanical interpretation. The denominator is the local stiffness margin, while the numerator is the net driving effect of shortening on the postural state. Thus, if shortening-induced forcing is strong or the local stiffness margin is small, the equilibrium becomes highly sensitive.

### 3.3 Second-order sensitivity and branch curvature

Differentiating once more gives

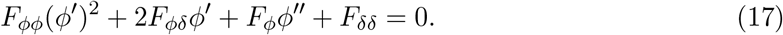

For the present model,

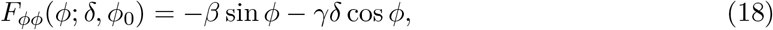

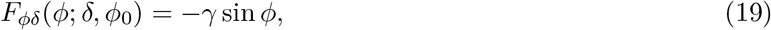

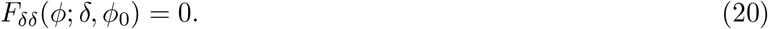

Hence,

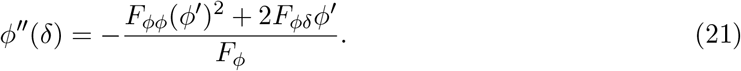

At *δ* = 0,

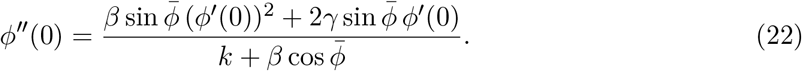

This quantity describes the local curvature of the equilibrium branch. In particular, if 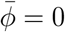, then sin 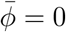 and

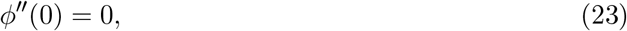

so the initial response is purely linear to second order. By contrast, baseline asymmetry immediately generates nonlinear curvature.

Thus the local expansion takes the form

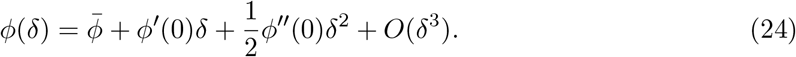

### 3.4 Local stability along a branch

An equilibrium along the branch is locally stable if and only if

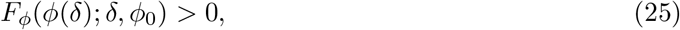

or equivalently,

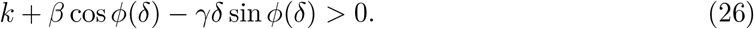

The key structural point is that the same quantity *F*_*ϕ*_ appears both in the stability criterion and in the denominator of the sensitivity formula:

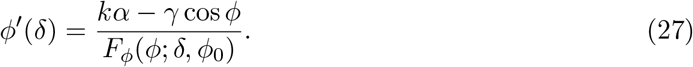

Therefore, unless the numerator vanishes,

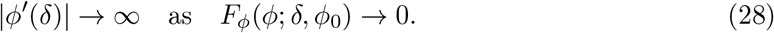

This means that the system becomes highly sensitive precisely when its local restoring margin becomes small. Clinically, even before stability is fully lost, a small change in shortening may already produce a large change in compensatory posture.

### 3.5 Degeneracy and onset of criticality

The nondegeneracy condition fails when

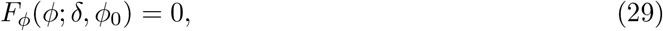

that is,

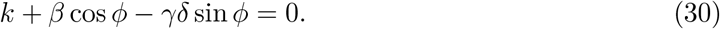

When this is imposed together with the equilibrium equation,

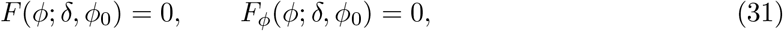

we reach the fold boundary. At such points, the equilibrium branch can no longer be represented as a regular graph, and qualitative branch geometry changes become possible.

### 3.6 Interpretation for fracture prognosis

The analysis above already has a clinically suggestive meaning. In ordinary parameter ranges, shortening induces a smooth compensatory response. But as the system approaches degeneracy, two things happen at once: branch sensitivity increases sharply, and the local stability margin shrinks. Together, these features create a mechanically fragile state in which small geometric differences may lead to disproportionately large functional consequences. This provides a natural explanation for why apparently similar fractures may have very different prognoses.

**Figure 2:**
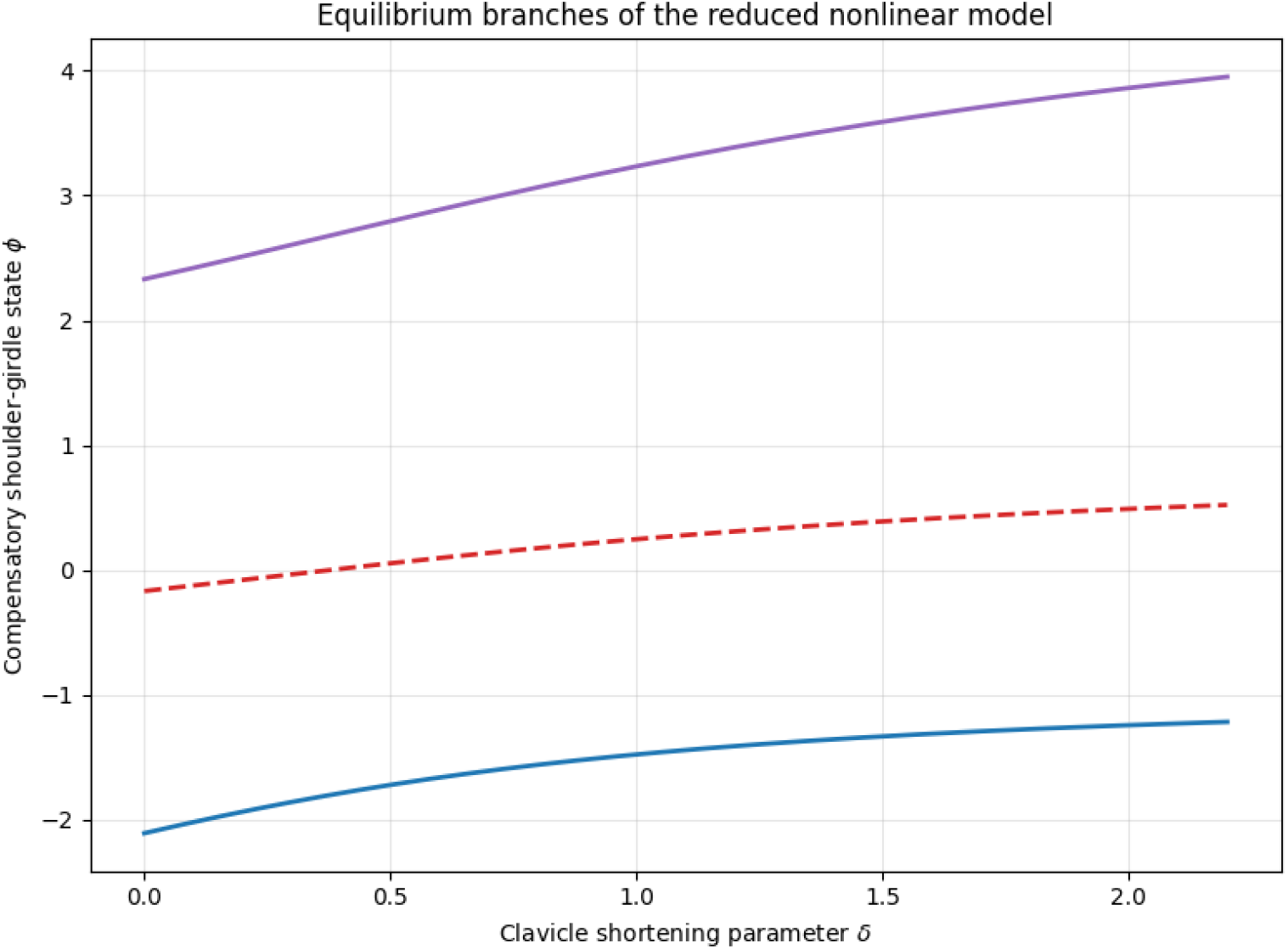
Representative equilibrium branches of the reduced model. Stable branches and unstable branches coexist, illustrating threshold-like response and mechanical fragility near criticality.

## 4. Fold and Cusp Structure

Section 3 identified the local precursor of instability. We now analyze the geometric structure produced when degeneracy actually occurs. We show that the simultaneous conditions *F* = 0 and *F*_*ϕ*_ = 0 yield fold bifurcation, and that under an additional condition the fold set itself becomes singular, producing a cusp. The resulting geometry gives a compact theoretical description of abrupt destabilization and strong patient-specific sensitivity.

### 4.1 Fold conditions

Recall the equilibrium equation

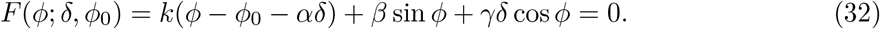

A fold point satisfies

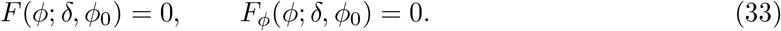

Since

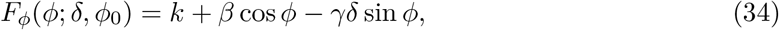

the fold system becomes

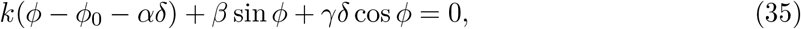

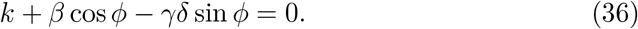

Whenever *γ* sin *ϕ* ≠ 0, we may solve for

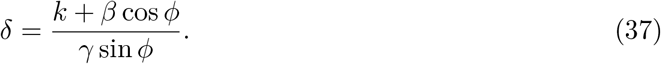

### 4.2 Local normal form of the fold

Let (*ϕ*^*^,*δ*^*^) be a fold point. Setting

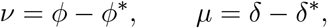

and expanding *F* near (*ϕ*^*^,*δ*^*^), we obtain

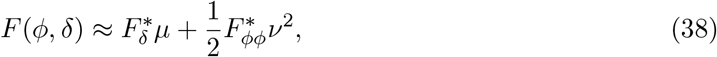

provided *F* (*ϕ*^*^,*δ*^*^) = 0 and *F*_*ϕ*_(*ϕ*^*^,*δ*^*^) = 0. Thus, if

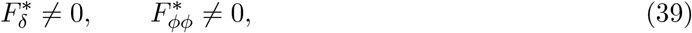

the equilibrium equation is locally equivalent to the standard fold normal form

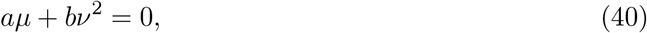

with *a, b≠* 0.

For the present model,

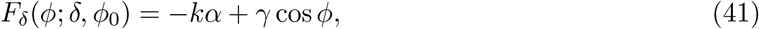

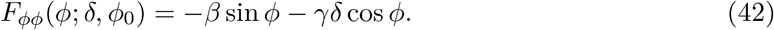

Under the corresponding nondegeneracy conditions, the equilibrium set has the square-root structure

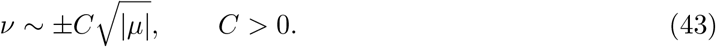

This means that two branches meet and terminate at the fold. Mechanically, one may interpret this as the collision and annihilation of a stable compensatory state with an unstable one. Beyond the fold, no nearby compensatory equilibrium exists.

### 4.3 Stable and unstable branches

From the stability criterion, a branch is locally stable if *F*_*ϕ*_ *>* 0 and unstable if *F*_*ϕ*_ *<* 0. Near a fold, the two branches produced by the local normal form therefore typically correspond to one stable branch and one unstable branch. In the energy picture, the stable branch corresponds to a local minimum, whereas the unstable branch corresponds to a local maximum. Thus, a fold is not merely a technical singularity; it marks the disappearance of a mechanically admissible compensatory state.

### 4.4 Parametric representation of the fold boundary

From equation (36), the fold boundary in parameter space can be written as

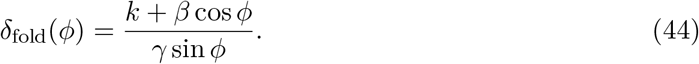

Substituting this into the equilibrium equation yields

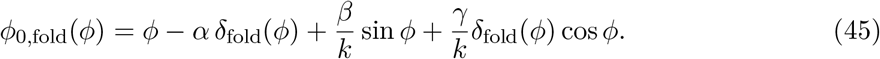

This representation makes clear that the critical boundary is generally not a single constant shortening threshold. Instead, it is a curved object in the (*δ, ϕ*_0_)-plane, shaped jointly by stiffness, geometric nonlinearity, and baseline posture.

### 4.5 Cusp conditions

A cusp occurs when the fold set itself becomes singular, which in this one-state-variable model corresponds to adding the condition

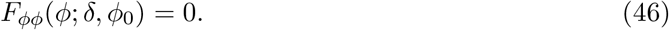

Thus, a cusp satisfies

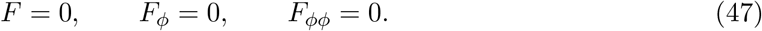

Using the explicit formulas, one obtains

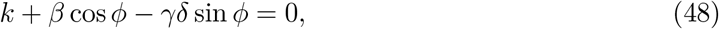

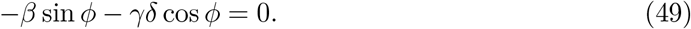

Squaring and adding these equations gives

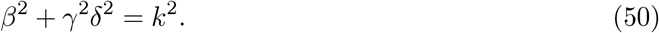

Hence, a real cusp exists if and only if

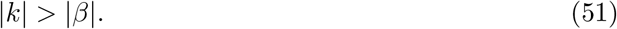

In that case,

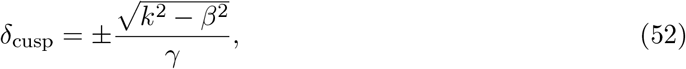

and

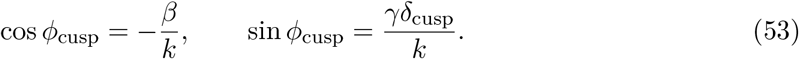

Substituting back gives

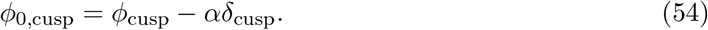

### 4.6 Nondegeneracy of the cusp

To verify that this is a genuine cusp rather than a higher-order accidental degeneracy, compute

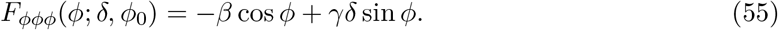

At the cusp point, this simplifies to

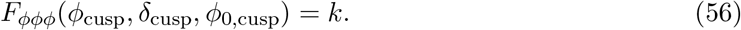

Therefore, provided

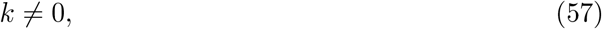

the cusp is nondegenerate in the standard sense.

### 4.7 Multistability and parameter sensitivity

Inside the cusp region, the equilibrium equation has three solutions, typically two stable and one unstable. Outside the cusp region, only one equilibrium remains. This gives a theoretical mechanism for pronounced inter-individual variability: two patients with similar fracture shortening may nevertheless lie in different regions of parameter space and therefore exhibit qualitatively different mechanical responses.

### 4.8 Clinical interpretation

Fold bifurcation explains why prognosis may deteriorate suddenly once a threshold is crossed. Cusp geometry explains why that threshold is not universal but depends on patient-specific baseline factors. In this sense, the jump threshold is not a fixed number but a section of a higher-dimensional critical surface.

**Figure 3:**
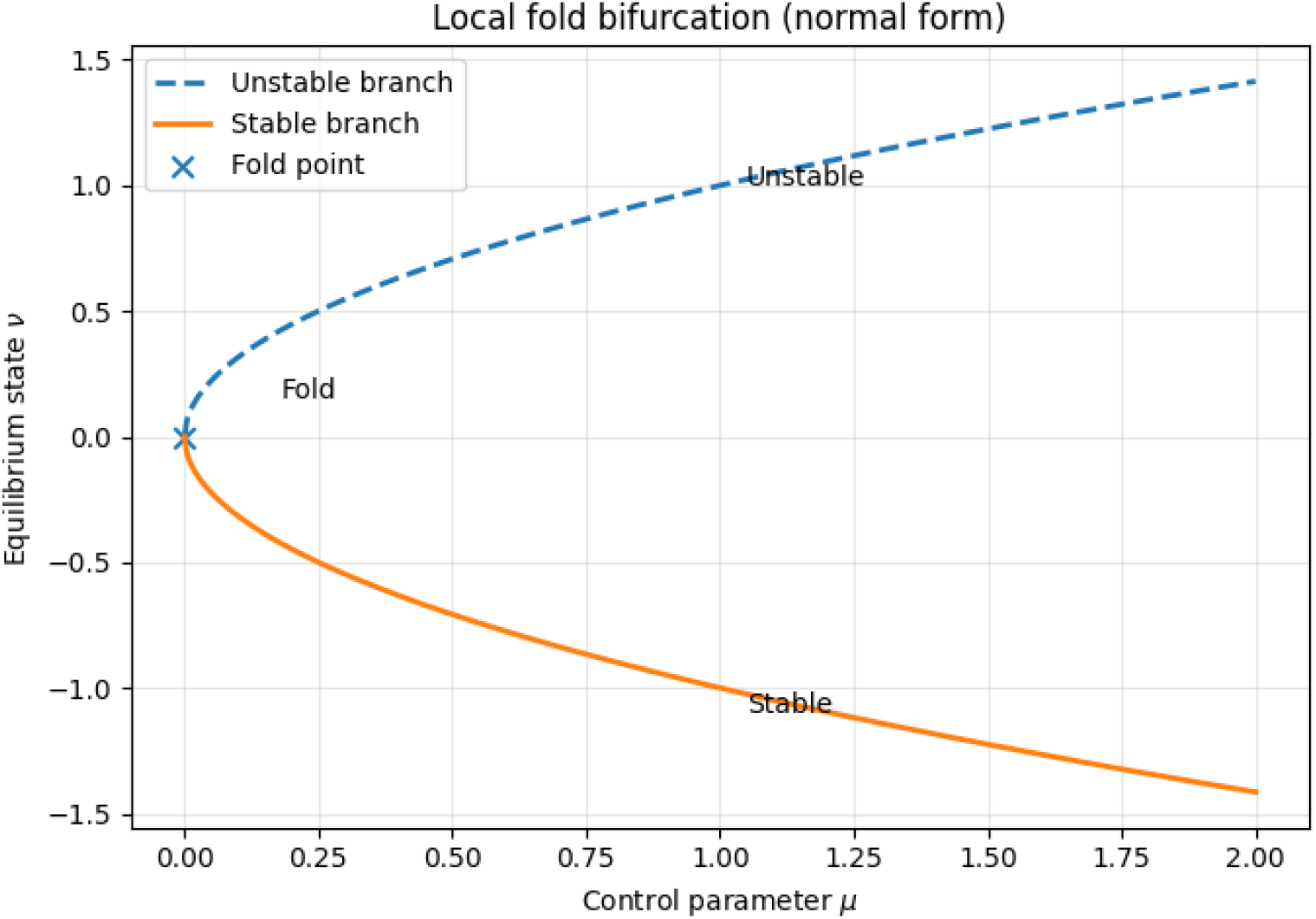
Representative fold-bifurcation geometry in the reduced model, showing the collision and disappearance of stable and unstable equilibria.

**Figure 4:**
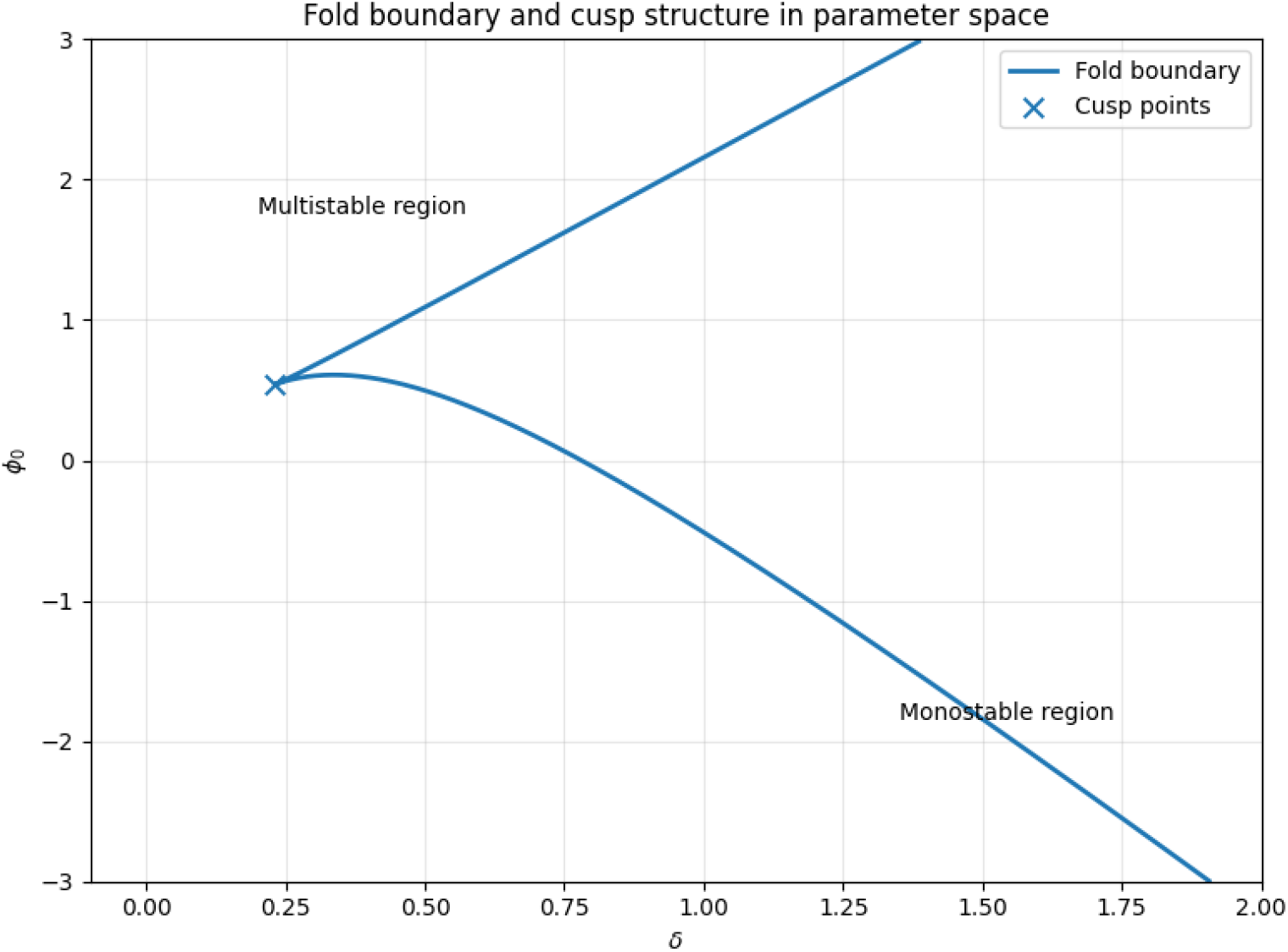
Cusp-organized parameter region in the (*δ, ϕ*_0_) plane. Inside the cusp region, multistability may occur; outside it, the system is typically monostable.

## 5. Optimization and Safety Margin

The bifurcation analysis shows that shortening may push the system toward a fold boundary, where stable compensation is lost. But treatment planning is not determined by stability alone. In practice, one must balance geometric correction against joint burden, muscular compensation, and fragility near instability. The goal of this section is to formalize that balance as an optimization problem defined on the stable equilibrium branch.

### 5.1 Optimization on the stable branch

Let

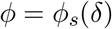

denote a stable equilibrium branch satisfying

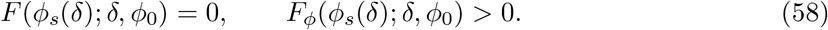

We then define the effective optimization problem

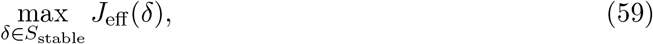

where

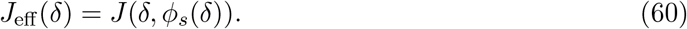

Thus, treatment choice is optimized not in the full state space, but along the mechanically admissible stable set.

### 5.2 A biomechanically motivated utility functional

We introduce

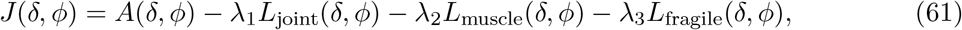

where *A* represents geometric or functional benefit, *L*_joint_ the cost of joint loading, *L*_muscle_ the cost of muscular compensation, and *L*_fragile_ a penalty for proximity to instability. The *λ*_*i*_ are weights.

The precise form of these terms is not unique and is not yet patient-calibrated. The purpose here is structural: any reasonable treatment objective should represent the fact that further correction may bring benefit, but also increases load, compensation cost, and fragility.

### 5.3 Effective derivative along the stable branch

Differentiating the effective utility gives

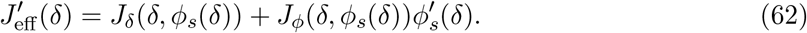

Using

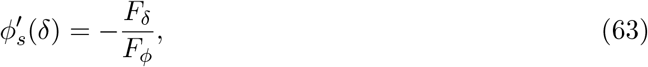

we obtain

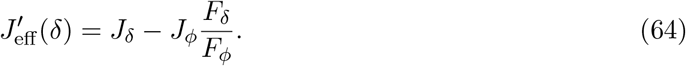

Hence an interior optimum satisfies

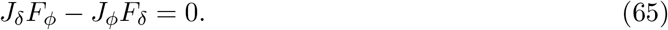

This identity reveals the key mechanism: even if direct benefit remains positive, the effective marginal cost can become very large when *F*_*ϕ*_ becomes small. In other words, approaching the fold amplifies fragility.

### 5.4 Why the optimum lies below the fold threshold

Let *δ*^*^ be the fold point at which the stable branch terminates. Near *δ*^*^, the stable branch has the local form

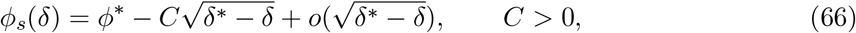

so that

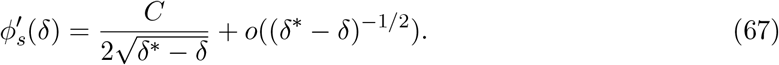

Meanwhile, the stiffness margin satisfies

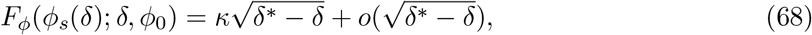

for some nonzero constant *κ*.

Thus, any reasonable fragility penalty diverges as *δ* → *δ*^*\*−*^, and even without an explicit singular term, the derivative formula already shows that effective cost becomes sharply unfavorable near the fold. Consequently, the optimum cannot generically occur at the fold boundary itself. Instead, one expects

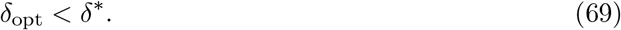

This means that the fold threshold is not an optimal target but only an upper bound of mechanical admissibility.

#### Proposition 5.1

(Optimal correction lies below the fold point). *Assume that the stable branch terminates at a nondegenerate fold point, and that the utility functional contains either an explicit fragility penalty or a nontrivial posture-related penalty that becomes unfavorable near the fold. Then the effective utility cannot generically attain a local maximum at the fold boundary. In generic situations, any local optimum satisfies*

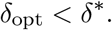

### 5.5 Definition of the safety margin

We therefore define the safety margin by

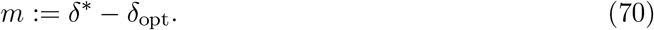

This quantity measures the distance between the optimal correction and the fold boundary in control-parameter space. If *m* is small, the chosen correction lies dangerously close to instability; if *m* is larger, the correction is more conservative but also more robust.

### 5.6 Local scaling law

Let

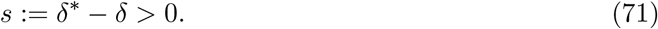

Near the fold, the effective utility may be expanded asymptotically as

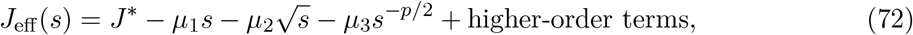

where the coefficients depend on the smooth benefit-cost terms and on the fragility penalty. This shows that the safety margin obeys a nontrivial scaling law governed by the competition between benefit and fragility. Stronger penalty on fragility increases the safety margin, whereas stronger benefit for additional correction decreases it.

### 5.7 Interpretation

The optimization framework leads to several conclusions. First, the maximal mechanically admissible correction and the optimal correction are distinct quantities. Second, the safety margin is not merely a heuristic buffer; it arises naturally from singular amplification near the fold. Third, the optimal correction is inherently patient-specific, since both the fold threshold and the effective utility depend on baseline parameters and clinical priorities.

## 6. Discussion and Clinical Implications

The central purpose of this paper is to provide a theoretical framework for understanding threshold-like deterioration in clavicle-fracture prognosis. Rather than treating shortening or displacement as purely linear predictors, we model the shoulder-girdle response as a reduced nonlinear mechanical system with equilibrium branches, stability boundaries, and bifurcation structures that depend on patient-specific baseline conditions.

### 6.1 What the model explains

One familiar clinical observation is that fractures that look similar on imaging can follow very different courses. Some are mechanically tolerated and heal satisfactorily, whereas others with similar apparent shortening develop persistent symptoms or poor healing. In the present model, shortening moves the system along a stable compensatory branch only up to a critical point. At the fold boundary, the stable equilibrium collides with an unstable one and disappears. This provides a mechanism for sudden loss of mechanical tolerance.

The cusp structure extends this explanation by showing that the critical boundary depends on baseline parameters. Thus, two patients with similar fracture geometry may still lie in different regions of parameter space and therefore have different prognoses. The model also explains why fragility may become clinically relevant before overt instability appears: as the fold is approached, the branch steepens and the effective stiffness margin shrinks, so the system becomes increasingly sensitive to small perturbations.

### 6.2 Reinterpreting empirical thresholds

This framework suggests a reinterpretation of empirical treatment thresholds. Clinically, surgeons often use rule-of-thumb shortening thresholds when deciding whether surgery should be considered. But the model indicates that such thresholds should not be understood as universal constants. Rather, they may be viewed as projections of a higher-dimensional critical surface involving baseline posture, joint geometry, soft-tissue state, and compensatory capacity.

The optimization analysis further distinguishes two clinically different quantities: the maximal limit of conservative treatment and the shortening level at which conservative treatment is actually optimal. This helps explain why cases lying very near a threshold may already be poor candidates for conservative management.

### 6.3 Limitations

The present model is intentionally simplified. It reduces the entire shoulder-girdle system to a single state variable and a small set of structural parameters. This is clearly insufficient for direct quantitative prediction in actual patients. Important anatomical features such as three-dimensional bone geometry, ligament arrangement, muscle-specific mechanics, scapulothoracic motion, and patient-specific loading are not explicitly represented.

In addition, the coefficients and functional forms have not yet been calibrated using clinical data. The model is therefore theoretical rather than predictive at this stage. It is also static: it does not incorporate bone biology, callus formation, vascular changes, or healing dynamics.

### 6.4 Why the simplified model is still useful

Despite these limitations, the model captures structural features that may remain important even in more realistic descriptions: coexistence of stable and unstable compensatory states, sudden loss of stable compensation at a fold bifurcation, cusp-organized inter-individual variability, and a safety margin below the instability threshold. These are not mere artifacts of overfitting, but generic consequences of nonlinear branch geometry.

For this reason, the present work can serve as a conceptual foundation for later anatomically richer or data-driven models. Even if more detailed finite-element or imaging-based approaches are developed, a reduced model such as this remains valuable as an organizing language for identifying the key mechanisms and clinically relevant observables.

### 6.5 Toward patient-specific prediction

A natural next step is to establish a mapping between the abstract structural parameters of the present model and measurable clinical quantities, such as radiographic shortening, clavicular angle, scapular posture, restriction of shoulder motion, or pain score. Only after such a mapping is built can the current framework be developed into a genuinely patient-specific predictive tool.

## Conclusion

We have proposed a minimal nonlinear biomechanical model for prognostic analysis of clavicle fractures. Within this framework, threshold-like deterioration arises naturally from fold bifurca-tion, patient-specific variability is organized by cusp geometry, and treatment planning can be formulated as an optimization problem on the stable equilibrium branch. The model predicts that the optimal safe correction lies strictly below the instability threshold, thereby generating a natural safety margin. Although simplified, the framework offers a mathematically coherent starting point for future patient-specific, data-calibrated, and anatomically richer models of clavicle-fracture prognosis.

